# A hierarchical regulatory cascade defines the temporal window of natural transformation in *Staphylococcus aureus*

**DOI:** 10.64898/2026.08.03.742410

**Authors:** Pierre Poirette, Shi Yuan Feng, Yolande Hauck, Yannis Arab, Sophie Quevillon-Cheruel, Stéphanie Marsin, Marine Rigot, Macha Dussouchaud, Roza Mohammedi, Thomas Hainaut, Paul Budisavljevic, Nicolas Mirouze

## Abstract

Natural transformation (NT), a major mechanism of horizontal gene transfer, promotes bacterial genome plasticity and contributes to the dissemination of antibiotic resistance. NT requires the development of competence, a transient and tightly regulated physiological state that is induced by environmental cues. Although competence has recently been demonstrated in the human pathogen *Staphylococcus aureus*, the regulatory architecture and temporal coordination governing its development remain poorly understood.

Using a highly sensitive luciferase-based transcriptional reporter, we demonstrate that NT gene expression in *S. aureus* is transient, thereby defining a temporal window during which DNA uptake and recombination can occur. We further identify two classes of NT genes with distinct regulatory requirements and activation kinetics: Class I genes require both SigH and ComK1 and are activated approximately 2.5 hours later than Class II genes, which are regulated exclusively by ComK1. Mechanistically, we demonstrate that ComK1 activates *sigH* transcription, establishing a hierarchical regulatory cascade that explains its essential role in Class I gene expression. Finally, time-resolved transcriptomic analyses identify the global regulator CodY as an early regulator of competence that differentially controls *comK1* and *sigH* transcription, thereby linking metabolic adaptation to competence development.

Together, our findings define the regulatory architecture that governs competence development in *S. aureus* and reveal how a hierarchical cascade integrates environmental and metabolic cues to restrict horizontal gene transfer to a precise developmental window. This work provides a mechanistic framework for understanding how bacteria coordinate physiological adaptation with genome diversification through NT.

**Author summary:** Bacteria constantly evolve by acquiring new genetic information from their environment. One important mechanism driving this process is natural transformation, which allows bacterial cells to take up and integrate exogenous DNA into their genomes. However, this process is not continuously active: bacteria must first enter a specialized physiological state called competence. How pathogens control the development of competence remain poorly understood.

In this study, we investigate how the human pathogen *Staphylococcus aureus* regulates natural transformation. Using a highly sensitive reporter system that allows us to monitor gene expression over time, we show that competence is activated only during a limited period, creating a defined window during which bacteria can acquire new DNA sequences. We uncover a hierarchical regulatory cascade involving the transcription factors ComK1 and SigH that controls the sequential activation of natural transformation genes. We further identify the metabolic regulator CodY as an upstream regulator that connects bacterial physiological state with the initiation of competence.

Our findings reveal that natural transformation is not a random or continuous process, but a precisely timed developmental program controlled by multiple regulatory layers. By defining how *S. aureus* coordinates environmental sensing, metabolism, and horizontal gene transfer, this work provides new insight into how bacterial pathogens generate genetic diversity and adapt to changing environments, including conditions that may promote the emergence of antibiotic resistance.

## Introduction

Natural transformation (NT), one of the three main mechanisms of horizontal gene transfer (HGT) in bacteria—alongside conjugation and transduction—enables the binding, internalization, and homologous recombination of exogenous DNA into the host chromosome (1–3). NT contributes to bacterial genomic plasticity, facilitating adaptation to fluctuating and stressful environments, and promoting the dissemination of antibiotic resistance as well as vaccine evasion (4). Unlike other HGT mechanisms, NT is entirely controlled by the recipient cell, which must enter a differentiated physiological state known as competence (5,6). Importantly, most transformable bacterial species do not constitutively express the proteins required for DNA binding, uptake, and recombination. Instead, competence development is typically transient, defining a limited window during which DNA internalization and transformation can occur (5,7).

The development of competence generally follows common principles. It is induced in response to environmental cues sensed by the recipient cell. These signals are integrated through complex signal transduction pathways that activate central competence regulators, which in turn control the expression of the “late” competence regulon, including all genes required for NT (8–12). Competence development can therefore be viewed as a succession of regulatory checkpoints that constrain the process within a defined temporal and physiological framework.

Despite conserved mechanisms for DNA uptake and processing, the regulation of competence has evolved independently across bacterial species, reflecting adaptation to diverse environmental conditions and organism-specific physiology (6,13). At each stage of competence development, species-specific regulators and regulatory mechanisms have been described. For instance, competence induction depends on a wide variety of environmental signals. While cell density and nutrient limitation are key triggers in *Bacillus subtilis* (14), sublethal concentrations of antibiotics or fever-like temperature can induce competence in *Streptococcus pneumoniae* (15,16). In response to such stimuli, signal transduction pathways modulate central competence regulators at multiple levels, including transcription, translation, and protein stability (5,6). Furthermore, the role of central competence regulator can be fulfilled by distinct types of proteins, such as alternative sigma factors, transcriptional activators, or co-regulators (5,6). This diversity highlights the substantial regulatory variation among transformable bacteria. For example, alternative sigma factors such as ComX in *S. pneumoniae* associate with RNA polymerase to direct transcription from specific promoter sequences characterized by distinct −10 and −35 elements (17). In contrast, the transcriptional activator ComK in *B. subtilis* is tightly controlled at both transcriptional and post-translational levels (14).

Competence for NT has only recently been demonstrated in the human pathogen *Staphylococcus aureus* (18). This discovery raises fundamental questions about how competence is regulated and deployed in a clinically important organism long thought to lack this capability. Recent work, including from our group, has begun to define the regulatory architecture underlying NT in *S. aureus*. Three central competence regulators have been identified: the alternative sigma factor SigH (19) (phylogenetically related to ComX in *S. pneumoniae*, (17)) and two transcriptional activators, ComK1 and ComK2 (20) (homologs to ComK from *B. subtilis*, (21)). Building on this framework, we developed a planktonic culture-based protocol that markedly increases the fraction of competent cells and enhances NT efficiency in *S. aureus*. Using this system, we showed that SigH and ComK1 are the primary regulators of NT genes expression (22) organizing the competence program into two distinct modules: Class I genes co-regulated by SigH and ComK1, and Class II genes controlled exclusively by ComK1 (22). Unfortunately, despite these advances, the temporal dynamics and higher-order regulatory logic governing competence activation in *S. aureus* remained largely unresolved.

Here, we confirm and extend these findings using *Photinus pyralis* luciferase as a sensitive transcriptional reporter in a new experimental framework, based on 96-well plates. We first demonstrate that competence in *S. aureus* is a transient and tightly regulated state, restricted to an approximately 10-hour window following oxygen limitation. Within this window, Class I and Class II genes exhibit distinct activation kinetics that mirror the dynamic expression patterns of *sigH* and *comK1*, revealing a temporally structured regulatory program. We further provide evidence that ComK1 directly or indirectly regulates *sigH* transcription, providing mechanistic insight into its essential role in coordinating Class I gene expression. Finally, time-resolved transcriptomic analyses identify candidate early regulators of competence and reveal a central role for the global regulator CodY in shaping the onset of competence by differentially modulating *sigH* and *comK1* expression.

Together, these findings establish competence in *S. aureus* as a transient, environmentally triggered, and hierarchically organized developmental program, with important implications for HGT and the evolution of this major human pathogen.

## Results

### Transient expression of NT genes defines a window of opportunity for HGT

As shown in **Fig. 1**, growth in sealed 96-weel plates, in CS2 medium under shaking led to rapid oxygen depletion, followed by a transient pause in growth, demonstrating that this experimental setup perfectly reproduced the competence-inducing conditions previously described in closed Falcon tubes (22). Under these conditions, expression from the *comG* operon promoter (P*_comG_*) displayed a sharp, peak-shaped profile. Induction occurred when oxygen levels reached their minimum (i.e. 0,3%), coinciding with the growth pause, and returned to basal levels approximately 10 hours later, upon entry into stationary phase (**Fig. 1**). A similar transient expression pattern was observed for all other NT genes tested (**Supp. Fig. 1a-f)**, with the exception of *recA*, which exhibited two distinct expression peaks (**Supp. Fig. 1g**).

**Fig. 1.**
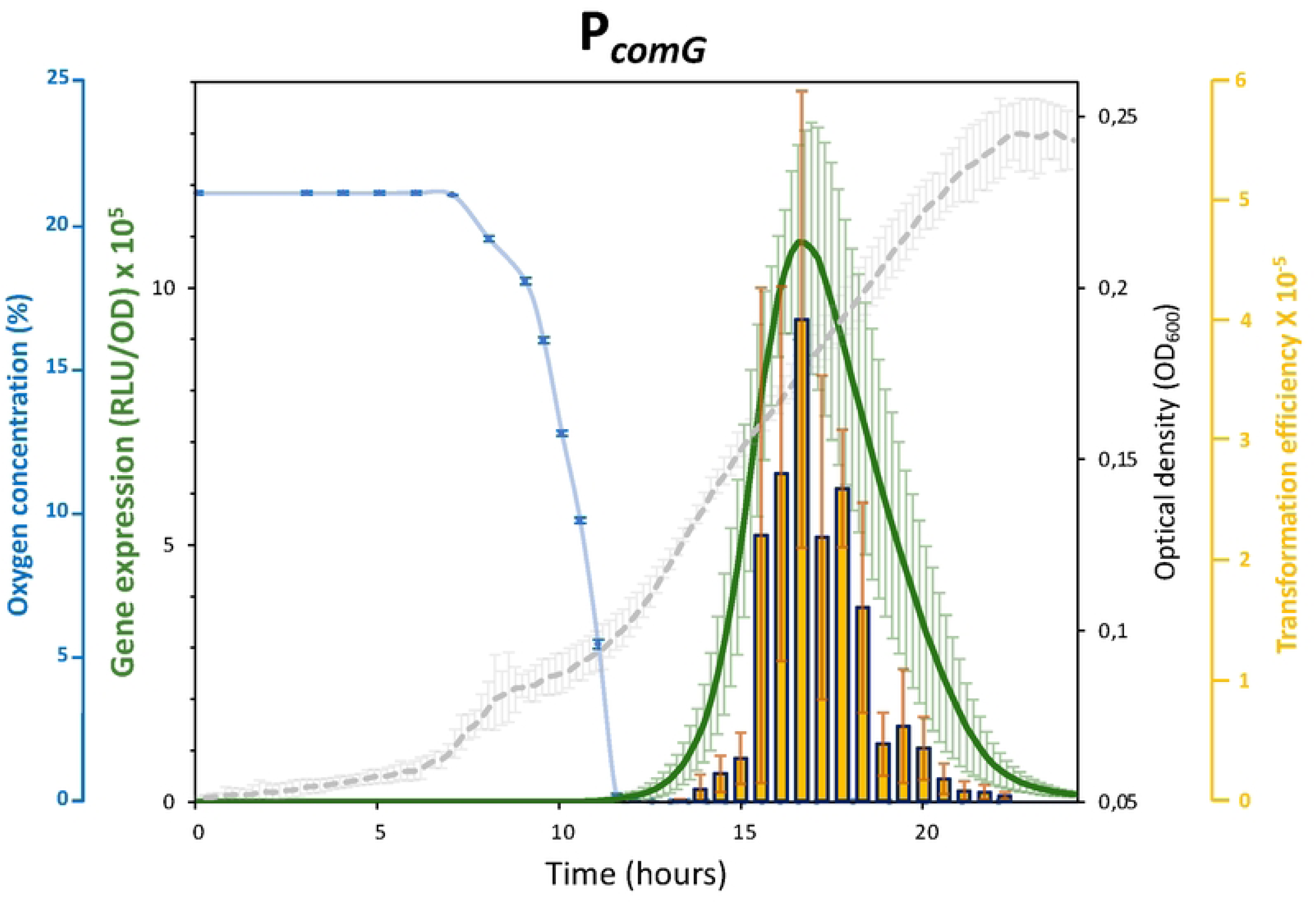
The competence transcriptional program is transient. Expression profile measured from the *comG* operon promoter (P*_comG_, St249*), expressed in Relative Light Unit (RLU) and normalized by the optical density (OD). The expression profile measured from the *comG* operon promoter (green curve) is shown as the mean of the transcription rate (with confidence intervals based on a Student Test; p-value<0.05, n=5) detected from 5 independent biological replicates (with at least 3 technical replicates each). The blue dotted line shows the evolution of the oxygen concentration in the culture (mean from 3 independent biological replicates with standard deviation) throughout growth (mean growth curve drawn in grey). The yellow bars represent the evolution of the transformation efficiency measured every 30 minutes (mean of 3 independent experiments with standard deviation; see more details in **Supp. Fig. 4**). The detection limit of the transformation efficiency was evaluated around 10^-8^.

Because oxygen concentration varies substantially during growth in CS2 medium, we assessed whether oxygen availability could influence luciferase activity. Although firefly luciferase has been reported to retain activity at extremely low oxygen levels (23), we experimentally confirmed that, in *S. aureus*, reporter activity remained detectable at oxygen concentrations at least tenfold lower (below 0.03 %, **Supp. Fig. 2**, (24)) than those measured under our experimental conditions (i.e. 0.3%). In addition, both the induction and decline phases of *P*_comG_ expression occurred while oxygen levels remained stable and low (i.e. 0.3%, **Supp. Fig. 3**), indicating that the observed transcriptional dynamics are not limited by oxygen availability.

**Fig. 2.**
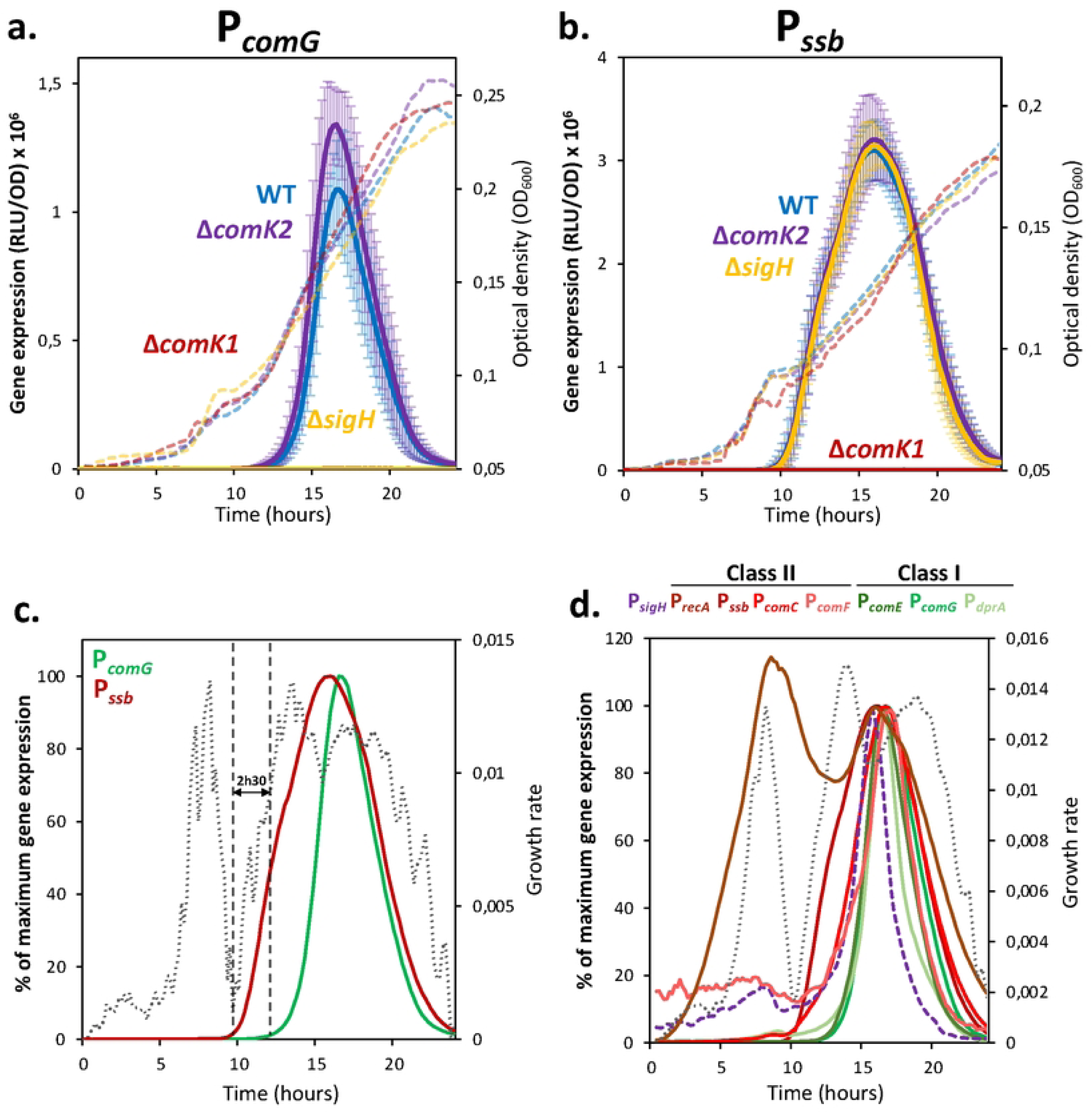
Two classes of natural transformation genes with specific expression timing. Comparison of the expression profiles (RLU/OD), shown as the mean of the transcription rate (with confidence intervals based on a Student Test; p-value<0.05, n=5), and measured from (a) the *comG* operon promoter (P*_comG_*), representing class I NT genes and the (b) *ssb* promoter (P*_ssb_*), a class II NT gene in WT (in blue, St249 and St254) and central competence regulator mutant strains (Δ*comK1* in red, St255 and St259; Δ*comK2* in purple, St443 and St260; and Δ*sigH* in yellow, St257 and St261). Expression profiles measured from P*_comG_* and P*_ssb_* (c, St249 and St2) or all the NT genes promoter (d) in a WT context, expressed in % of their maximum transcription rate (P*_sigH_*, St264; P*_recA_*, St377; P*_ssb_*, St254; P*_comC_*, St406; P*_comF_*, St508; P*_comE_*, St390; P_comG_, St249 and P*_dprA_*, St474). The mean growth rate (grey dotted line) is also represented as a temporal reference (see **Supp. Fig. 4** for details). In panel (c), a 2,5 hours delay between the initiation of expression from P*_ssb_* and P*_comG_* is materialized by vertical dotted lines.

**Fig. 3.**
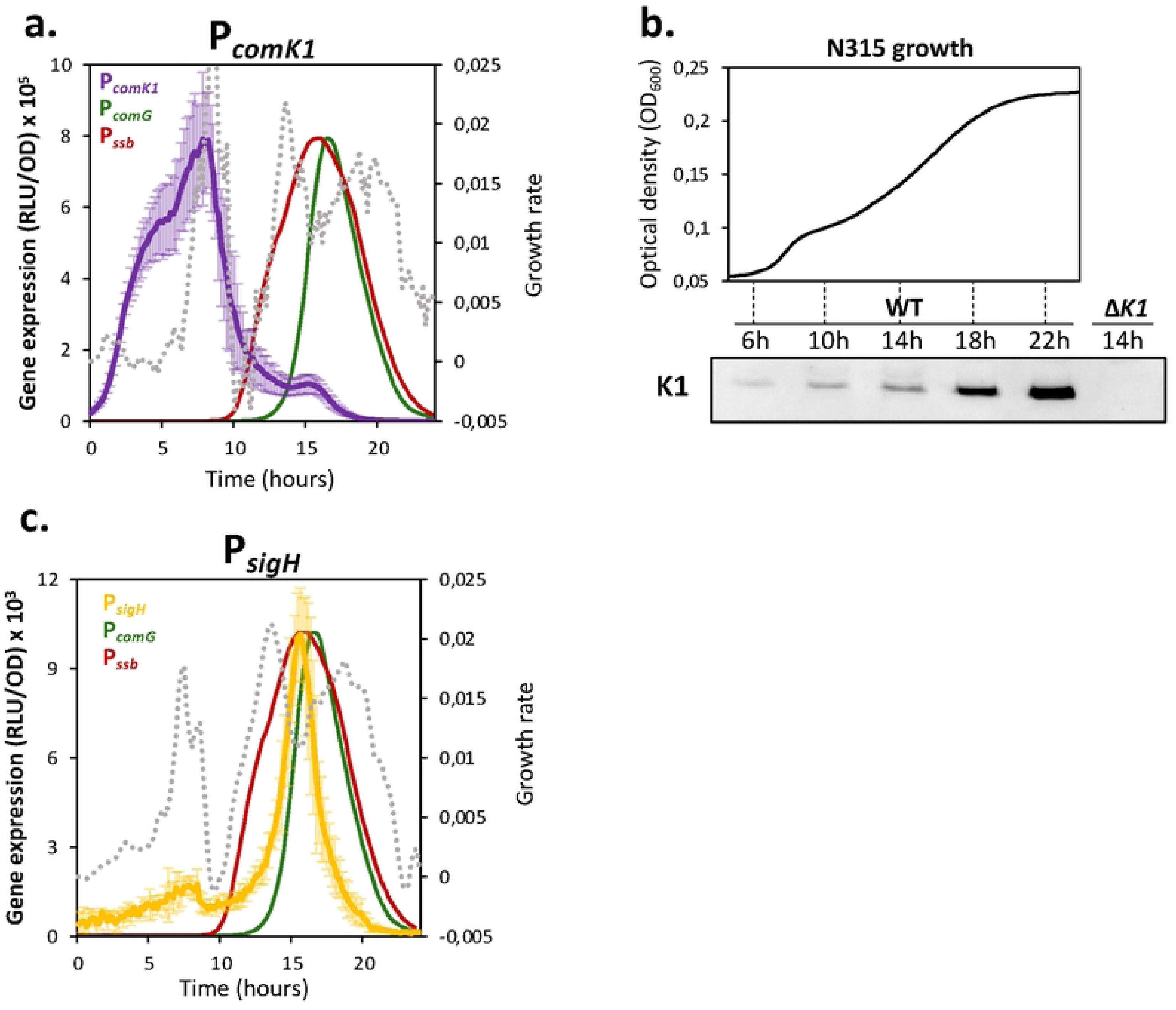
Expression of central competence regulators controls the NT genes timing. (a) Expression profile (RLU/OD), shown as the mean of the transcription rate (with confidence intervals based on a Student Test; p-value<0.05, n=5), and measured from the *comK1* promoter (P*_comK1_*, St389 in purple) along growth. P*_comG_* (St249, in green) and P*_ssb_* (St254, in red) expression profiles are presented as temporal references (their maximum expression level is adjusted to that of *comK1*). The mean growth rate (grey dotted line) is also represented as a temporal reference (see **Supp. Fig. 5a** for details). (b) Quantification of ComK1 protein by Western blot in a WT strain, St389 (6h, 10h, 14h, 18h, 22h) as well as in a Δ*comK1* mutant strain, St416 (14h). Growth (OD_600_) of a WT strain is shown in top panel to visualize at which growth phase each western blot sample corresponds. (c) Expression profile (RLU/OD) measured from the *sigH* promoter (P*_sigH_*, St264, in yellow) along growth. P*_comG_* (St249, in green) and P*_ssb_* (St254, in red) expression profiles are presented as temporal references (their maximum expression level is adjusted to that of P*_sigH_*). The mean growth rate (grey dotted line) is also represented as a temporal reference (see **Supp. Fig. 5b** for details).

We next verified if the transient expression of the NT genes defines a corresponding window for HGT in *S. aureus*. By measuring the transformation efficiency of a wild type strain, every 30 minutes along growth in CS2 medium, we demonstrated that the apparition of transformants closely coincided with the onset of *comG* expression (P*_comG_*), that the peak of transformants matched the peak of *comG* expression and that no transformants were detected once expression returned to basal levels (**Fig. 1 and Supp. Fig. 4**). This tight temporal coupling indicates that competence in *S. aureus* is restricted to a narrow physiological window and underscores the requirement for potential regulatory control to enable transient NT.

**Fig. 4.**
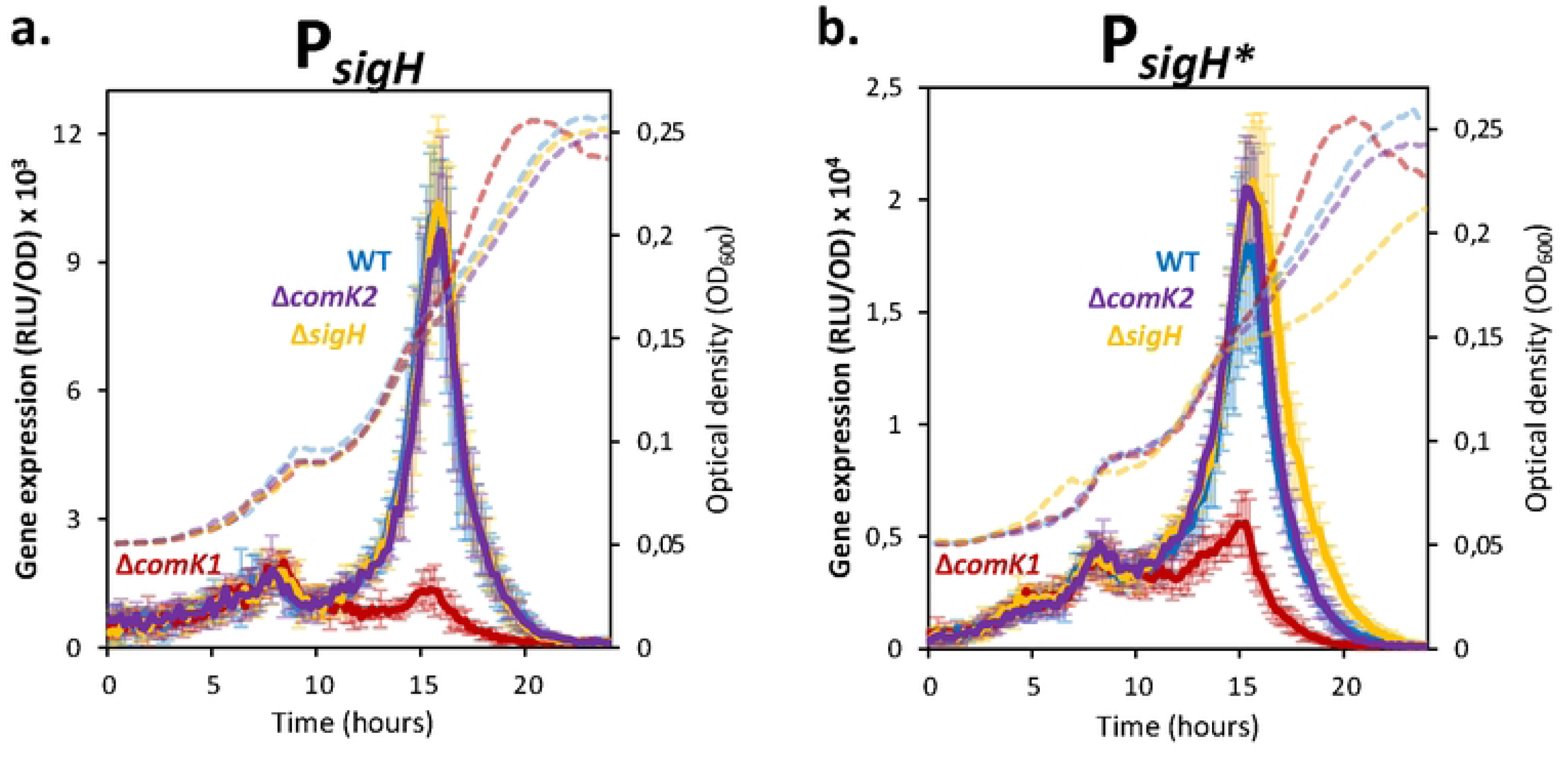
ComK1 activates *sigH* transcription. P*_sigH_* (a) and P*_sigH*_* (b) expression profiles (RLU/OD), shown as the mean of the transcription rate (with confidence intervals based on a Student Test; p-value<0.05, n=5), and measured in WT (St264 and St289) and central competence regulator mutant strains (Δ*comK1*, St415 and St414; Δ*comK2*, St426 and St423 and Δ*sigH*, St422 and St418). Growth (OD600) of the WT (blue), Δ*comK1* (red), Δ*comK2* (purple) and Δ*sigH* (yellow) mutant strains harboring pRIT-P*_sigH_*-luc or pRIT-P*_sigH*_*-luc strains are represented as dotted lines.

### Two classes of NT genes differentially regulated

Highlighting the complexity of competence regulation in *S. aureus*, we previously identified two classes of NT genes defined by distinct regulatory requirements: Class I genes, which require both SigH and ComK1, and Class II genes, which depend solely on ComK1 (22). To validate this classification, we compared the expression of representative Class I (P*_comG_*) and Class II (P*_ssb_*) promoters in wild-type and mutant strains lacking their respective key competence regulators (**Fig. 2a** and **2b**). As expected, P*_comG_* activity was abolished in both *ΔsigH* and *ΔcomK1* mutant strains (**Fig. 2a**) whereas P*_ssb_* expression was specifically dependent on ComK1 and unaffected by the absence of SigH (**Fig. 2b**).

Next, we extended this analysis to additional NT genes and confirmed their assignment to each class when already known (*comG*, *ssb*, *comC* and *comF*, (22)) and identified their class when not yet investigated (*comE*, *recA*, *dprA*) (**Supp. Fig. 5**). Ultimately, the *comG* and *comE* operons, as well as *dprA*, were classified as Class I genes, whereas *ssb*, *comC*, *recA*, and the *comF* operon belonged to Class II. Notably, a subset of genes (*dprA*, *comC*, and *comF*) retained residual expression in the absence of either *comK1* or *sigH* (**Supp. Fig. 5c, 5e** and **5f**), suggesting that additional regulatory inputs, revealed by the high sensibility of the luciferase reporter, contribute to their transcription. In the case of *recA*, only the second expression peak was dependent on ComK1 (**Supp. Fig. 5g**), while the first peak persisted independently of competence, consistent with its role during vegetative growth (25).

**Fig. 5.**
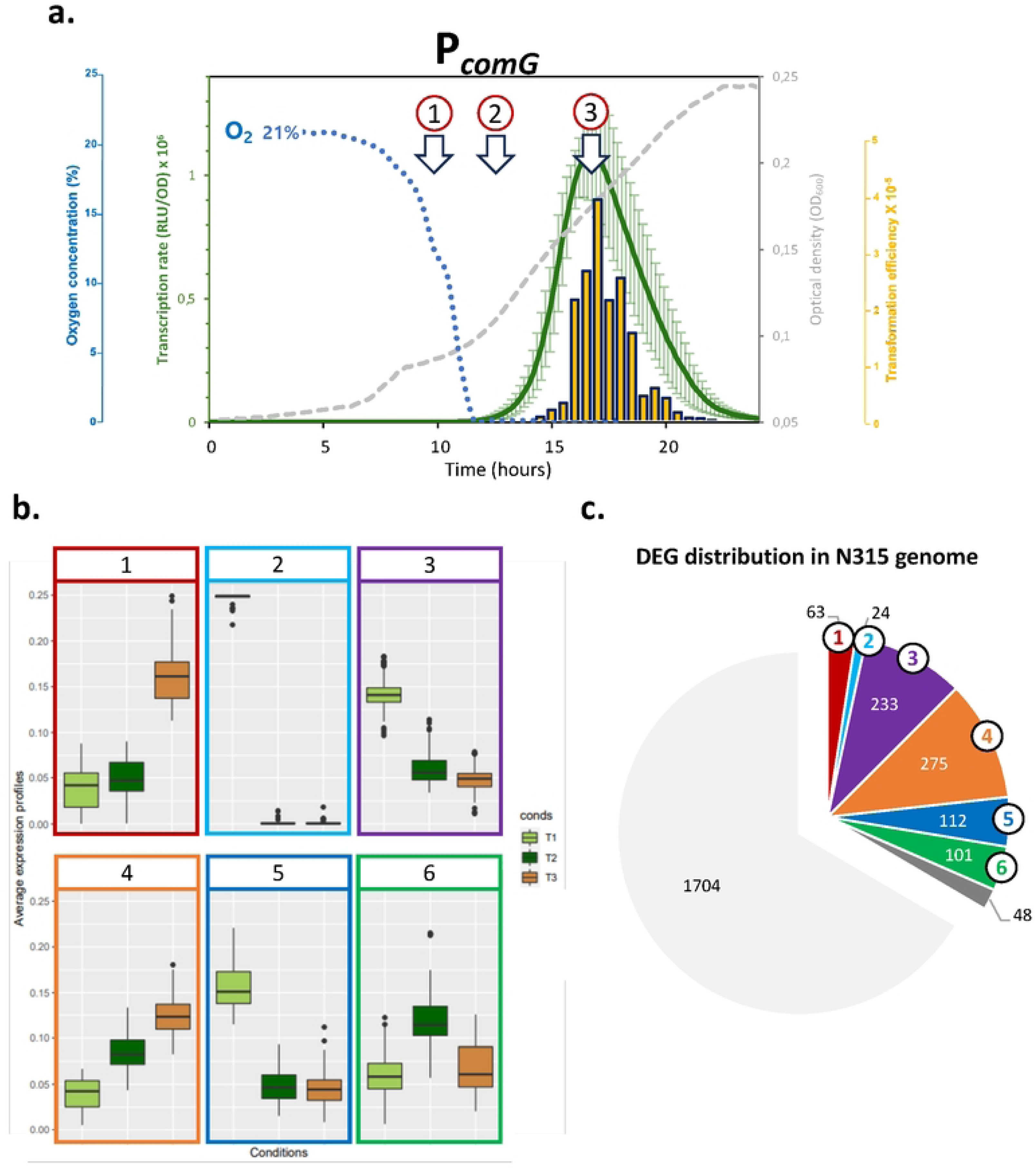
Clusters of differentially expressed genes. (a) RNA-seq samples are taken at different time points through the growth corresponding to (T1) the first early time for which the oxygen concentration was still around 12%, then (T2) when the oxygen concentration is reaching 0.2% right before the expression of the NT genes and finally (T3) when the expression of the NT reaches its maximum. (b) Average expression profiles of the 6 gene clusters from 4 RNA-seq technical replicates showing T1 (light green), T2 (dark green) and T3 (orange) and (c) their proportion among the N315 genome. Genes in light grey are not DEGs and genes in dark grey were DEGs not associated to any cluster.

### Two classes of NT genes differentially expressed through time

The high reproducibility of growth and reporter activity across strains enabled a detailed comparison of the temporal dynamics of NT gene expression (**Fig. 2c** and **2d**). Growth in CS2 medium is characterized by a transient pause during early exponential phase, likely reflecting adaptation to anaerobic metabolism, as illustrated by a sharp decrease in growth rate between 8 and 10 hours (**Supp. Fig. 6**). To account for minor variations in growth between strains, we used the growth rate as a temporal reference to align expression profiles. This analysis revealed a consistent delay of approximately 2.5 hours between the onset of Class II gene expression (e.g., *ssb*) and that of Class I genes (e.g., *comG*) (**Fig. 2c**). This temporal offset was conserved across all NT genes examined, with expression initiation clustering according to class (**Fig. 2d**). These findings indicate that competence gene expression follows a temporally ordered program and suggest that the distinct activation kinetics of Class I and Class II genes are intrinsically linked to their regulatory architecture.

**Fig. 6.**
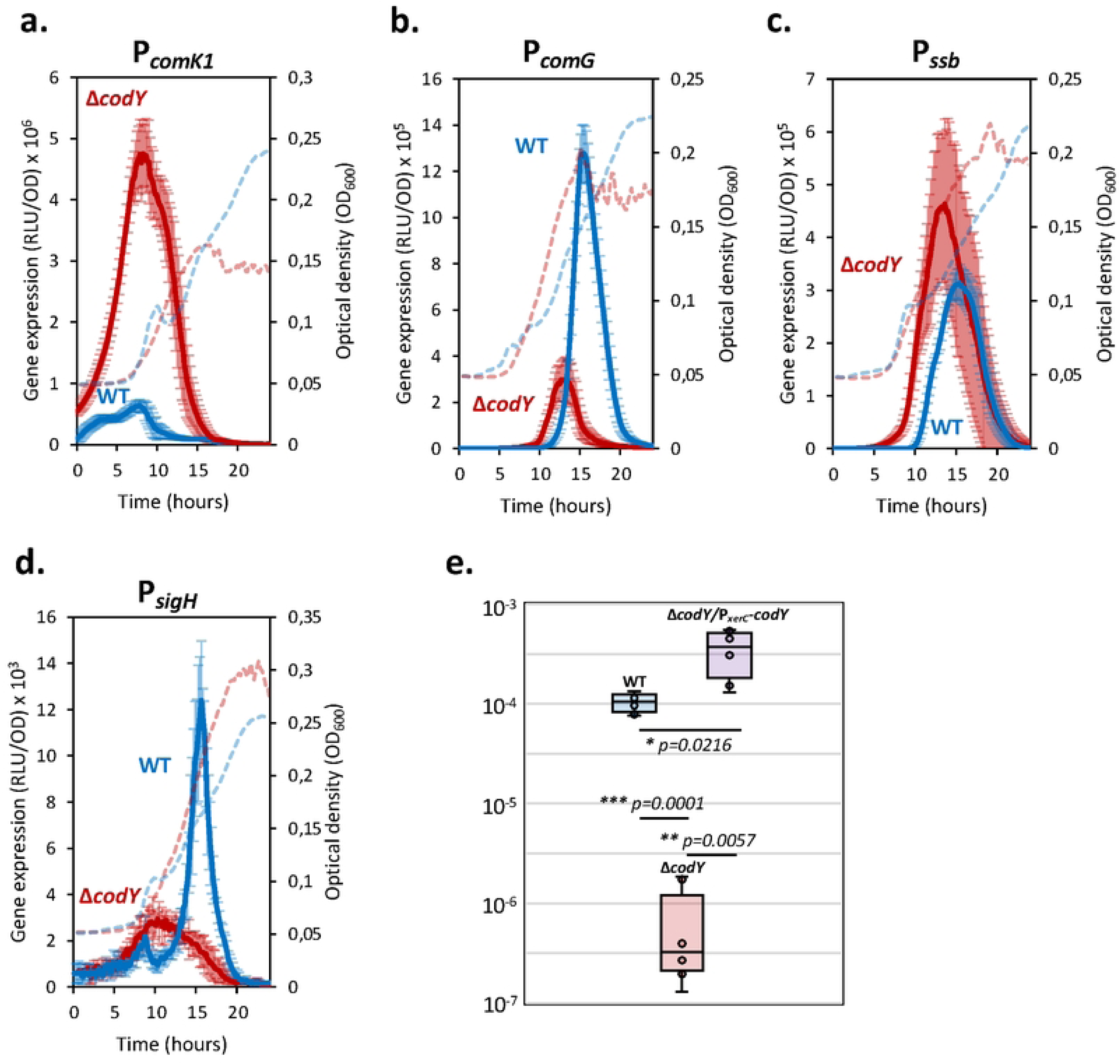
CodY is a major competence regulator. Expression profiles (RLU/OD), shown as the mean of the transcription rate (with confidence intervals based on a Student Test; p-value<0.05, n=5), and measured from the *comK1* (P*_comK1_*, a), *comG* (P*_comG_*, b), *ssb* (P*_ssb_*, c) and *sigH* (P*_sigH_*, d) promoters are compared in WT (St389, St249, St254 and St264 respectively, in blue) and a Δ*codY* (St393, St519, St392 and St457, respectively in red) strains. The growth curves of the WT and Δ*codY* mutant strain (harboring their respective luciferase fusions) are respectively shown as blue and red dotted lines. Boxplot of the maximal transformation efficiency (panel e.) of a WT (St12), a Δ*codY* (St388) and Δ*codY* pCN34-P*xerC-codY* (St766) strain. A Fisher test and a Welch test were performed on n=6 replicates. *:p-value <0.05; *:p-value<0.01; ***:p-value <0.001.

### Central competence regulators activation

We next asked whether the temporal offset between Class I and Class II gene expression could be explained by the expression dynamics of the two central competence regulators, *sigH* and *comK1*. To address this, we constructed luciferase transcriptional fusions to the promoter of both genes (**Fig. 3** and **Supp. Fig. 7**).

**Fig. 7.**
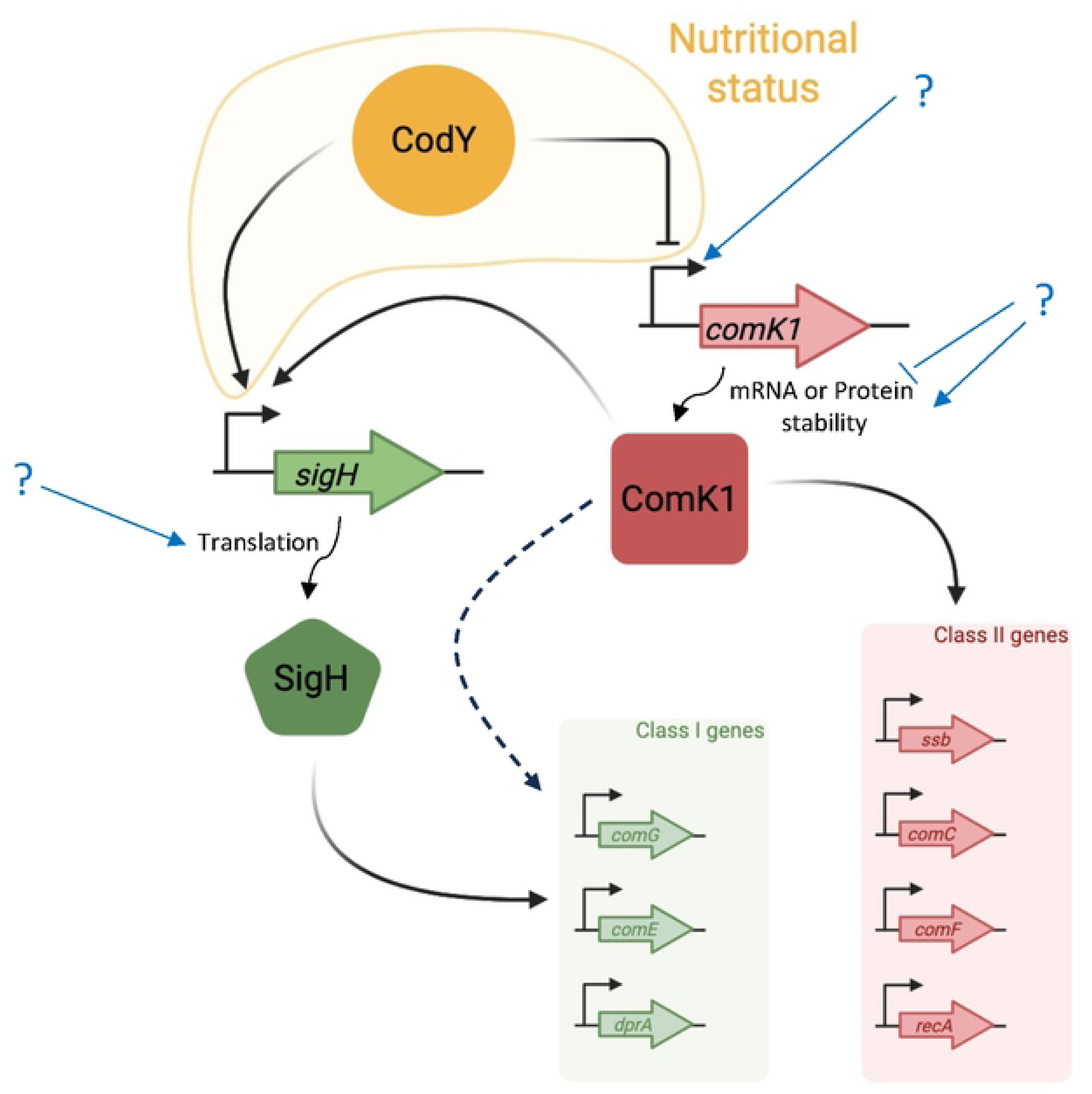
Model for the central regulation of competence in *S. aureus*. The two central competence regulators ComK1 and SigH are both essential for NT by controlling the expression of two sets of NT genes. Class I genes are controlled by SigH and ComK1 while class II genes are exclusively activated by ComK1. However, since we demonstrated that ComK1 activates sigH transcription, ComK1 essentiality for Class I genes expression is, at least in part, due to the activation of sigH by ComK1 (the potential direct control of Class I genes by ComK1 is materialized by a dashed arrow). Here, we also showed that CodY, a global transcriptional regulator diagnosing the nutritional status of the cell, is also a major competence regulator in *S. aureus*, that inhibits comK1 transcription early during growth and later activates sigH transcription. Finally, additional regulatory pathways still need to be discovered and characterized (blue arrows). The stimuli and regulators activating comK1 transcription remain unknown while we propose that comK1 mRNA or ComK1 protein stability need to be modulated. In parallel, we also know that sigH translation represents an addition layer of regulation to modulate the development of competence in *S. aureus*^15^.

Expression from the *comK1* promoter (P*_comK1_*) was detectable from the earliest time point (**Fig. 3a**), likely reflecting residual expression from the BHI pre-culture (**Supp. Fig. 8**). P*_comK1_* expression increased rapidly, reaching a maximum around 8 hours of growth before declining. A second, smaller peak was observed at later time points, around 15 hours of growth, representing ∼12% of the initial maximum (**Fig. 3a**). Notably, P*_comK1_* expression was relatively low at the time of maximal NT gene expression, prompting us to examine ComK1 protein levels by Western blot (**Fig. 3b** and **Supp. Fig. 9**). ComK1 protein was already detectable early during growth (around 6 hours), consistent with its transcriptional profile, but accumulated progressively throughout competence development. Importantly, ComK1 levels continued to increase even as NT gene expression declined (**Fig. 3a** and **3b**).

In contrast, *sigH* promoter (P*_sigH_*) expression remained low during the first half of the growth curve and increased sharply after the growth pause, reaching a maximum at approximately 16 hours (**Fig. 3c**). Given that post-transcriptional regulation mediated by the *sigH* 5ʹ untranslated region (5ʹ-UTR) has been proposed as a key control point (18), we generated a modified reporter (P*_sigH*_*) in which the repeated and inverted sequences lying on *sigH* SD sequence were replaced by *sigA* ribosome binding site. This modification did not alter the overall expression dynamics but resulted in a twofold increase in signal intensity (**Supp. Fig. 10**), consistent with partial translational repression mediated by the native 5ʹ-UTR, as previously suggested (18).

Strikingly, the onset of the main P*_sigH_* expression peak coincided with the initiation of Class II gene expression, exemplified by *ssb*, which is exclusively regulated by ComK1 (**Fig. 3c** and **Fig. 2d**). This temporal correlation led us to hypothesize that *sigH* transcription may be controlled, directly or indirectly, by ComK1.

### ComK1 activates *sigH* transcription

We next assessed the contribution of the central competence regulators SigH, ComK1, and ComK2 to the expression of *comK1* and *sigH*. Deletion of these regulators revealed that P*_comK1_* expression was modestly repressed in the absence of SigH (i.e. less than 2-fold, **Supp. Fig. 11**). In contrast, P*_sigH_* expression was strongly dependent on ComK1. Deletion of *comK1* resulted in an approximately 10-fold reduction in P*_sigH_* promoter activity, whereas deletion of *sigH* or *comK2* had no significant effect (**Fig. 4**). Notably, residual P*_sigH_* expression persisted even in the absence of ComK1, suggesting that additional regulatory pathways contribute to its activation. Importantly, the impact of *comK1* deletion was similar for both the native P*_sigH_* and the modified P*_sigH*_* (**Fig. 4a** and **4b**), indicating that ComK1 primarily influences *sigH* at the transcriptional level rather than through post-transcriptional mechanisms. Together, these results demonstrate that ComK1 directly or indirectly activates *sigH* transcription, providing a mechanistic explanation for the requirement of ComK1 in the expression of Class I NT genes.

### Time-series transcriptomic for competence onset analysis

To identify early regulators acting upstream of the central competence factors, we performed time-series transcriptomic profiling, a powerful approach to capture dynamic biological processes. By comparing global gene expression across defined time points during growth, we aimed to identify differentially expressed genes (DEGs) and coordinated expression patterns associated with the onset of competence in *S. aureus*.

Transcriptomic analysis was conducted on a wild-type strain in four biological replicates at three key time points (**Fig. 5a**): T1, corresponding to early exponential phase with oxygen levels around 12%; T2, at which oxygen levels decreased to ∼0.2%, immediately preceding NT gene expression; and T3, corresponding to maximal NT gene expression. Differential expression and co-regulation analyses were performed using DicoExpress (27), identifying 856 DEGs (|log₂FC| > 1) that grouped into six clusters based on their expression profiles at different time points (**Fig. 5b, 5c and Supp. Fig. 12**).

As an internal validation, we first examined gene categories previously shown to respond to oxygen limitation in CS2 medium (22), including NT genes, anaerobic metabolism pathways, and virulence-associated genes. As expected, NT genes were predominantly expressed at T3, with 10 out of 14 genes clustering in group 1 (hereafter referred to as the NT cluster) (**Supp. Fig. 13**). *dprA* was assigned to cluster 4, consistent with its early competence-independent expression at T2 (compare **Supp. Fig. 5c** and **Supp. Fig. 13**), while *comEA* (and the *comE* operon) fell just below the differential expression threshold. Unexpectedly, *comK1* also clustered with NT genes, despite its earlier expression peak (**Fig. 3a**). This apparent discrepancy may reflect high transcript stability and progressive accumulation of ComK1 protein, as suggested by our Western blot analyses (**Fig. 3b**).

We next examined genes involved in anaerobic metabolism. Consistent with expectations, they were strongly upregulated at T2 compared to T1 and were primarily distributed across clusters 4 and 6 (**Supp. Fig. 14**). Genes in cluster 4 displayed sustained expression between T2 and T3, suggesting a role in maintaining anaerobic metabolism, whereas genes in cluster 6 showed transient induction, likely reflecting an acute response to oxygen depletion.

Finally, in agreement with previous observations (22), many virulence-associated genes were downregulated under oxygen-limited conditions (**Supp. Fig. 15**). These genes were distributed across clusters 2, 3, 5, and 6 and exhibited decreased expression at T3.

### Identification of new potential early competence regulators

We next leveraged the time-series transcriptomic dataset to identify candidate early regulators acting upstream of the late competence cascade. To this end, we focused on a curated list of 124 transcription factors and alternative sigma factors encoded in *S. aureus* N315 genome (26). Among the DEGs, we identified 34 transcriptional regulators associated with a wide range of cellular functions (**Supp. Table 1**), highlighting a broad regulatory rewiring during growth in CS2 medium. These included regulators linked to competence (*comK1*), metabolism (*pdxR*, *fapR*, *gntR* family members), virulence (*rot*), and biofilm formation (members of the *sar* family). We also detected several two-component system response regulators involved in stress adaptation, including *phoP*, *kdpE*, *nreC*, *vraR*, and *hptR*.

Importantly, transcriptional regulators that do not meet stringent DEG thresholds may still exert substantial regulatory effects on downstream networks. This is exemplified by *sigH*, which does not appear as a DEG despite strong induction of its regulon, underscoring the sensitivity of regulatory networks to modest changes in regulator abundance or activity. A similar case was observed for *codY*, encoding a major pleiotropic transcriptional regulator (27), which displayed a log₂(FC) of 0.93 between T1 and T2, just below the cutoff used for clustering. Notably, 69% of genes belonging to the CodY regulon were identified as DEGs in our dataset (**Supp. Table 2**), indicating substantial remodeling of CodY-controlled pathways during competence onset and development. Given that CodY has previously been shown to regulate *comK* expression in *B. subtilis* (28), these observations prompted us to investigate whether CodY also plays a role in the early stages of competence regulation in *S. aureus*.

### CodY is a major competence regulator in *S. aureus*

To investigate the role of CodY in the early stages of competence development in *S. aureus*, we first compared *comK1* expression in wild-type and Δ*codY* strains during growth in CS2 medium (P*_comK1_*, **Fig. 6a**). Strikingly, *comK1* expression increased more than sevenfold in the absence of CodY, indicating that CodY acts as a repressor of *comK1* during competence onset (**Fig. 6a**). To confirm that this phenotype was not due to polar effects of the gene deletion, we complemented the Δ*codY* strain by ectopic expression of *codY*, which partially restored *comK1* wild type expression levels (**Supp. Fig. 16a**). Given that CodY has been shown in *B. subtilis* to directly regulate *comK* expression through promoter binding (28), we next searched for CodY consensus binding motifs in the *comK1* promoter region of *S. aureus*. Two putative CodY binding sites were identified, supporting the possibility of a conserved direct regulatory interaction (**Supp. Fig. 17**).

We then assessed the downstream consequences of CodY inactivation on NT gene expression. Consistent with increased ComK1 activity, expression of *ssb* (P*_ssb_*, a Class II gene exclusively regulated by ComK1) was significantly increased in the Δ*codY* strain, although to a lesser extent than *comK1* itself (**Fig. 6b**). This phenotype was fully reversed upon genetic complementation (**Supp. Fig. 16b**). In contrast, expression of the *comG* operon (P*_comG_*, a Class I gene requiring both ComK1 and SigH) was significantly reduced in the Δ*codY* strain (**Fig. 6c**), despite the expected increase in ComK1 levels. Complementation of *codY* also restored *comG* expression to wild-type levels (**Supp. Fig. 16c**).

This apparent discrepancy suggests that CodY exerts opposing effects on the two central competence regulators. While increased ComK1 levels in the Δ*codY* background would be expected to enhance *sigH* activation and consequently promote Class I gene expression, the observed repression of *comG* indicates that *sigH* may itself be negatively affected in the absence of CodY. To test this hypothesis, we analyzed *sigH* transcription in the Δ*codY* strain. Strikingly, *sigH* expression was significantly reduced in the absence of CodY (**Fig. 6d**), indicating that CodY positively influences *sigH* transcription. This effect was partially restored upon complementation (**Supp. Fig. 16d**). Consistent with a potential direct regulatory role, we also identified a putative CodY binding site within the *sigH* promoter region (**Supp. Fig. 17**).

Finally, given the global and opposing effects of CodY on key competence regulators and NT genes, we evaluated its functional impact on NT. The Δ*codY* strain exhibited a dramatic reduction in transformation efficiency (∼1000-fold compared to wild type; **Fig. 6e**), consistent with the strong decrease in *comG* expression. As expected, ectopic expression of *codY* restored transformation efficiency to wild-type levels (**Fig. 6e**). Together, these results identify CodY as a central early regulator of competence in *S. aureus*, acting as a bidirectional transcriptional modulator of *comK1* and *sigH* and thereby integrating metabolic status with the transient activation of NT (**Fig. 7**).

## Discussion

In most naturally transformable species, with the notable exception of certain Gram-negative bacteria such as *Neisseria spp.* and *Helicobacter pylori*, competence is a transient differentiation state. Here, we demonstrate that *S. aureus* follows this general principle. NT gene expression is restricted to a ∼10-hour window under the planktonic conditions tested, thereby limiting HGT to a defined temporal period. Expression of NT genes follows a peak-shaped profile, initiated upon oxygen depletion and subsequently shut off by yet-unidentified mechanisms. The competence window in *S. aureus* is substantially longer than that reported for *S. pneumoniae* (20–40 min, (11,12)), but is comparable to that observed in *B. subtilis* (4–5 hours, (29)). This extended window in *S. aureus* may reflect its slower growth under oxygen-limited conditions in CS2 medium, thereby ensuring that a sufficient fraction of the population can enter competence.

We previously defined two classes of NT genes based on regulatory control (22). Here, we confirmed and refined this classification: Class I genes (*comG*, *comE*, *dprA*) require both SigH and ComK1, whereas Class II genes (*comF*, *comC*, *recA*, *ssb*) depend exclusively on ComK1. Our high-sensitivity luciferase reporter further revealed residual expression of several genes, suggesting additional layers of regulation or competence-independent transcriptional inputs. These observations point to a more complex regulatory architecture than previously appreciated, potentially involving additional regulators acting in parallel or upstream of the central competence pathway.

A key finding of this study is the existence of a temporal delay between the two classes of NT genes, with Class II genes activated approximately 2.5 hours before Class I genes. Importantly, this temporal organization does not reflect the order of functional steps in DNA uptake, as genes encoding the DNA transport machinery (e.g. *comG* and *comE*) are expressed later. Rather, we show that this temporal hierarchy is explained by the expression dynamics of the central regulators: *comK1* is expressed early, whereas *sigH* expression peaks later and coincides with the onset of NT genes expression. Moreover, we demonstrate that *sigH* transcription is positively regulated by ComK1, providing a mechanistic basis for the delayed activation of Class I genes. This regulatory cascade likely explains the essential role of ComK1 in Class I gene expression, although a direct contribution of ComK1 to Class I promoters cannot be excluded (**Fig. 7**).

The regulatory organization uncovered here shows striking parallels with the competence network of *Vibrio cholerae*, where TfoX regulates QstR to control downstream gene classes (30). Similarly, both organisms exhibit a modular architecture composed of multiple central regulators controlling distinct subsets of NT genes. Such multilayered regulation likely enables fine-tuning of competence in response to environmental fluctuations. In *S. aureus*, additional complexity arises from incomplete repression of *sigH* in a *comK1* mutant, post-transcriptional regulation of SigH (**Fig. 7**, (18)), and yet-unknown signals governing *comK1* induction as well as NT genes shutdown. Furthermore, the apparent disconnect between *comK1* luciferase transcriptional profile and the ComK1 protein abundance suggests the involvement of additional regulatory layers, potentially affecting *comK1* mRNA stability or translation, or ComK1 protein stability and activity (e.g., protein–protein interactions or post-translational modifications). Altogether, these observations collectively indicate that competence is controlled by a highly interconnected regulatory network integrating multiple environmental and intracellular cues.

Time-resolved transcriptomics further refined this model by identifying six clusters of co-regulated genes, capturing global transcriptional reprogramming during growth in CS2 medium. While most NT genes clustered together, discrepancies between RNA-seq and luciferase-based profiles highlight the complementary nature of these approaches. Luciferase reporters capture promoter activity with high temporal resolution and can partially reflect post-transcriptional regulation, whereas RNA-seq integrates mRNA abundance, including stability effects, at specific time points. For example, the early luciferase signal for *comK1* likely reflects rapid transcriptional activation followed by mRNA stabilization, whereas the absence of *sigH* from DEG lists likely reflects its low transcript abundance despite strong regulatory impact.

Beyond competence genes, transcriptomics revealed a broader adaptive response involving metabolic adaptation, anaerobic metabolism, stress responses, and virulence regulation. Among identified transcription factors, CodY emerged as a central early regulator integrating metabolic status with competence onset and development. We demonstrate that CodY represses *comK1* while promoting *sigH* expression, thereby exerting opposite effects on the two central competence regulators. This dual role positions CodY as a key regulatory node at the interface between metabolism and developmental reprogramming. Similar nutrient-dependent control of competence has been described in *B. subtilis* (28) and *Lactococcus lactis* (31), suggesting evolutionary conservation of CodY-mediated regulation across Gram-positive bacteria.

Importantly, the opposing effects of CodY on *comK1* and *sigH* expression could reveal an important regulatory crossroad at the heart of competence onset (**Fig. 7**). By simultaneously modulating both central regulators, CodY likely acts as a buffering system that prevents premature or excessive activation of competence. We propose that CodY integrates metabolic signals to finely tune the onset and amplitude of the competence program. Ultimately, the phenotypes revealed along competence development (i.e. transcriptional regulations, transformation efficiency) perfectly qualifies CodY as a major early competence regulator in *S. aureus*. Future work will be required to determine whether CodY acts directly on both promoters and to identify the environmental cues controlling this regulatory axis.

Together, our results support a model in which competence in *S. aureus* is a transient, hierarchically organized developmental program controlled by sequential regulatory layers (**Fig. 7**). Early metabolic adaptation through CodY-dependent regulatory rewiring, controls expression of both *sigH* and *comK1*, which in turn induce downstream NT genes. This cascade generates a defined temporal window of competence, ensuring efficient yet tightly controlled HGT events.

## Materials and Methods

### Bacterial strains and growth conditions

All bacterial strains and plasmids used in this study are listed in **Supp. Table 3**. *S. aureus* strains were grown at 37°C under aerobic conditions in Brain Heart Infusion (BHI), Competence-inducing Synthetic medium (CS2) or GS medium. CS2 and GS media were freshly prepared from stock solutions as previously described (18).

*E. coli* IM08B or DH10B strains were used as vector hosts to amplify plasmids and were cultured in Luria-Bertani (LB) medium at 37°C under aerobic conditions. When needed, one or several of the following antibiotics were added to the culture media depending on the bacterial genotype: 100 µg/mL ampicillin (Amp), 10 µg/mL chloramphenicol (Cm), 200 µg/mL kanamycin (Kan) and 50 µg/mL neomycin (Neo).

### Plasmids construction and transformations

Promoters of interest were inserted in the pRIT plasmid, upstream to the luciferase gene (32), using the Gibson assembly method. The promoters, the gene encoding the luciferase and the pRIT were first amplified with dedicated primers. All the oligonucleotides used in this study are listed in **Supp. Table 4**. In every primer’s tail, we designed 25–50 bp homologous overlap regions between a given fragment and the adjacent fragment. The PCR were performed using the Phusion High-Fidelity DNA Polymerase (purchased from Thermo Scientific). All the PCR fragments were then purified using a standard commercial silicon column (PCR cleanup kit, Macherey- Nagen) and verified by gel electrophoresis for the absence of non-specific or minor PCR fragments.

Gibson assembly master mix was prepared by adding 320 μL of 5× ISO buffer (25% w/v PEG-8000; 500mM Tris-HCl, pH 7.5; 50mM MgCl2; 50mM DTT; 5 mM NAD; 1mM each dNTP), 0.64 μL 10 U/μL T5 exonuclease, 20 μL of 2 U/μL Phusion polymerase, 160 μL of 40 U/μL Taq DNA ligase and 699.36 μL of water (all reagents were purchased from New England Biolabs). Five microliters of the two DNA fragments mixture containing 100 ng of linear pRIT plasmid and a 3-fold excess of inserts were added to 15 μL of Gibson assembly master mix. The reaction tubes were then incubated at 50 °C for 1 hr. Finally, 1 μL of the assembly reaction was transformed into IM08B electro-competent *Escherichia coli* cells. Transformed *E. coli* cells were incubated at 37 °C on LB agar with 100 μg/mL ampicillin. In order to select the transformants containing the expected plasmid, colony PCR was performed. The resulting plasmids were extracted from positive transformants (overnight cultures) and purified using a commercial kit (NucleoSpin Plasmid extraction kit, Macherey-Nagen). All the plasmids were verified by sequencing (GATC company). Finally, in order to obtain the final reporter strains, *S. aureus* electrocompetent cells were transformed with each constructed plasmid.

### Construction of *S. aureus* deletion mutants

In order to investigate the role of CodY, allelic replacement constructs were cloned into the temperature sensitive pIMAY plasmid (33) as explained in (22). Briefly, fragments corresponding to 1 kbp flanking regions of the *codY* gene were amplified from *S. aureus* N315 genomic DNA using primers flanked by restriction enzyme sites compatible with the multiple cloning site (MCS) of pIMAY. All the primers used for cloning in the present study are listed in **Supp. Table 4**. The resulting plasmids were electroporated into the IM08B *E. coli* strain, verified by colony PCR, purified and sequenced. *S. aureus* N315ex woϕ strain was finally transformed by electroporation. After selection of pIMAY integration into the chromosome, and subsequent exit, the resulting clones were screened by colony PCR to identify clones containing the desired mutation. Mutant strains were finally verified by PCR and DNA sequencing.

### Construction of codY complemented strain

In order to confirm that the phenotypes observed in the *codY* mutant strain were due to the absence of *codY*, we constructed a plasmid (pCN34-P*_xerC_*-*codY*) to ectopically express *codY* under the control of the promoter found upstream from the operon where *codY* is found in fourth place (Note that pCN34 and pRIT have compatible origins of replication so that the two plasmids can coexist in the same cells). Briefly, fragments corresponding to 1 kbp upstream of the *xerC* gene (first gene in the *codY* operon), the *codY* gene and the pCN34 plasmid were amplified using primers compatible with the Gibson assembly method. All the primers used for cloning in the present study are listed in **Supp. Table 4**. In every primer’s tail, we designed 25–50 bp homologous overlap regions between a given fragment and the adjacent fragment. The PCR were performed using the Phusion High-Fidelity DNA Polymerase (purchased from Thermo Scientific). All the PCR fragments were then purified using a standard commercial silicon column (PCR cleanup kit, Macherey- Nagen) and verified by gel electrophoresis for the absence of non-specific or minor PCR fragments.

Gibson assembly master mix was prepared by adding 320 μL of 5× ISO buffer (25% w/v PEG-8000; 500mM Tris-HCl, pH 7.5; 50mM MgCl2; 50mM DTT; 5 mM NAD; 1mM each dNTP), 0.64 μL 10 U/μL T5 exonuclease, 20 μL of 2 U/μL Phusion polymerase, 160 μL of 40 U/μL Taq DNA ligase and 699.36 μL of water (all reagents were purchased from New England Biolabs). Five microliters of the DNA fragments mixture containing 100 ng of linear pCN34 plasmid and a 3-fold excess of inserts were added to 15 μL of Gibson assembly master mix. The reaction tubes were then incubated at 50 °C for 1 hr. Finally, 1 μL of the assembly reaction was transformed into IM08B electro-competent *Escherichia coli* cells. Transformed *E. coli* cells were incubated at 37 °C on LB agar with 100 μg/mL ampicillin. In order to select the transformants containing the expected plasmid, colony PCR was performed. The resulting plasmids were extracted from positive transformants (overnight cultures) and purified using a commercial kit (NucleoSpin Plasmid extraction kit, Macherey-Nagen). All the plasmids were verified by sequencing (GATC company). Finally, the *S. aureus codY* mutant strain was transformed by electroporation. The complemented strain was finally verified by PCR and DNA sequencing (*codY* gene deletion and presence of pCN34-P*_xerC_*-*codY*).

### Luciferase assay

For the detection of luciferase activity, we followed a similar protocol as published in (32). Briefly, cells were first isolated from −80 °C stock on the BHI plate. Three colonies were inoculated in 5 mL of BHI and incubated at 37 °C with shaking at 180 rpm until OD reached 2.5. This pre-culture was then centrifuged, washed in fresh CS2 medium, and used to inoculate 1 mL of fresh CS2 medium (OD = 0.5) in 2 mL Ependorf tubes. Then, serial 10-fold dilutions were performed in 2 mL Eppendorf tubes. 50 μL of the 10^-2^ dilution culture were finally distributed in each of three wells in a 96-well black plate (PerkinElemer). 2.5 μL of luciferin were added to each well to reach a final concentration of 1.5 mg/ml (4.7 mM). The plate was then filmed (Viewseal #676070, Greiner Bio One) to prevent gases exchanges and recreate the conditions previously published (22). The cultures were incubated at 37°C with double orbital shaking at 180 rpm in an Envision Xcite Multilabel Reader (PerkinElmer) equipped with an enhanced sensitivity photomultiplier for luminometry. The temperature of the top of the plate was maintained at 39°C to avoid condensation. Relative Luminescence Unit (RLU) and OD 600 were measured at 10 min intervals. Note that OD measurements are realized in the plate reader through 50 μL of medium which provide a light pathway significantly lower than in cuvette with 1 cm path length. Expression profiles are corrected by the number of cells present in the well at each point (RLU/OD). Each expression profile is the mean of the OD-corrected transcription rate from 5 independent biological replicates, themselves means of at least 3 technical replicates. The error bars are confidence intervals, based on a Student Test (p-value<0.05; n=5).

For some experiments, in order to compare the temporal dynamic of genes expression between strains, we used the growth rate to ensure that the key growth features (i.e. the growth pause occurring after oxygen depletion), were concomitant in each culture.

### Oxygen concentration measurements

Oxygen concentrations were measured using the SP-PSt3-SA23-D3-OIW oxygen sensor spots (PreSens GmbH, Regensburg, Germany). These sensor spots were attached at the bottom of a well (from a 96- well plate). The plate was then filmed (Viewseal #676070, Greiner Bio One) to prevent gases exchanges and recreate the conditions previously published (22). The sensor spots are covered with an oxygen-sensitive coating where molecular oxygen quenches the luminescence of an inert metal porphyrine complex immobilized in an oxygen-permeable matrix. This process guarantees a high temporal resolution and a measurement without drift or oxygen consumption. The photoluminescence lifetime of the luminophore within the sensor spot was measured using a polymer optical fiber linked to an oxygen Meter (Fibox 4 trace; PreSens GmbH). Excitation light (505 nm) was supplied by a glass fiber, which also transported the emitted fluorescence signal (600 nm) back to the oxygen meter.

Briefly, an oxygen measurement was realized, through the plastic bottom of the well (from the 96-well plate), by simply approaching the optical fiber from the sensor spot. At each time point, the oxygen concentration was measured three times and the results provided represent the mean of these three measurements. In our experiments, the oxygen concentration was measured every 30 min. The experiment shown in **Fig. 1** have been repeated 3 times. The oxygen detection limit of the sensors used is around 0.03% (which is 10-time higher than the lowest oxygen concentration detected in CS2 medium).

### Natural transformation of competent *S. aureus* cells

Cells were naturally transformed following the protocol previously published (22). Briefly, at each time point (every half hour), 100 μL of cells were harvested from the 96-well plate, centrifuged at 10,000g for 1 min at 4 °C, resuspended in 100 μL of fresh CS2. 500 ng of chromosomal donor-DNA (associated to a chloramphenicol resistance) was added and incubated at 37 °C for 2.5 h with agitation at 180 rpm (a “no DNA” control was performed at each experiment). 10 or 90 μL where finally mixed with 25 mL of melted BHI agar pre-cooled to 55 °C together with an antibiotic, and the mixture was poured into Petri dishes. After solidification, the plates were incubated at 37 °C for 48 h. At each time point, the viability was also evaluated by serial dilution on BHI agar plates. Transformation efficiencies were finally calculated by dividing the number of transformants detected in 1mL of culture by the total number of cells in the same volume. The transformants were also verified by PCR.

The experiments shown in **Fig. 1 and Fig. 6e** have been repeated 3 times (see **Supp. Fig. 2** for details of Fig. 1) to provide strong statistical relevance. Our detection limit in all the natural transformation experiment is evaluated around 10^-8^.

### ComK1 purification and Preparation of ComK1 antibodies

A His6 tagged version of ComK1 was expressed in Escherichia coli BL21 Gold using the pET29a vector. Protein expression was induced in 800mL of cultures by using 2xYT medium supplemented with 800µL of IPTG (100µg/mL). After an overnight incubation at 15°C, cultures were collected and the pellet resuspended in 80mL of Phosphate Buffer (20mM Phosphate+1M NaCl, pH5.6). After a night at -20°C, to induce cell lysis, the cultures are sonicated (Branson Sonifier 250, 4 cycles of 50 sec, pulse at 90%, intensity at 40 arbitrary units). The cytoplasmic fraction was purified by centrifugation (18,000 × g, 30min, 20 °C). The supernatant was then loaded into a Ni-NTA resin. Elutions obtained at 200mM and 400mM of Imidazole were then concentrated using ultrafiltration by centrifugation (Vivaspin protein concentrator spin columns, Cytiva). These samples were then further purified using an ÄKTA pure™ size-exclusion chromatography system. Chosen fractions containing the purified protein were then finally concentrated using ultrafitration, aliquoted in Eppendorf tubes, frozen in liquid nitrogen and conserved at -80°C.

ComK1 antibodies. 1mg of purified ComK1-His protein was used to immunize a rabbit on day 1 and sample the final bleed on day 63 (Davids Biotechnologie GMBH). Finally, the antiserum was further purified using an antigen-specific affinity purification (Davids Biotechnologie GMBH).

### Western-blot analysis

After being grown in 10mL of CS2 medium in Falcon 50 tubes at 37°C with shaking at 180 rpm, ∼2.108 cells were collected punctually through growth by quick centrifugation at 10000 rpm for 2 minutes and were frozen in liquid nitrogen. For total protein extracts, the cell pellets were diluted in 100 μL of Laemmli sample buffer and denatured for 5 minutes at 95°C. 10 μL of protein extracts were then separated on a 16% polyacrylamide gel at 320V, 70mA for 35 min. Another migration was run on a gel that was subsequently stained with Coomassie blue and washed with acetic acid to control the homogeneity of the sample loading. The proteins were then transferred onto a 0.2μm nitrocellulose membrane (AmershamTM Protran^TM^) at 150V, 300mA for 1h30 min. The membrane was blocked in a TBS-Tween1X-milk3% solution for 1h at 4°C with shaking. After multiple washes, the membrane incubated with 0.01mg/mL affinity-purified anti-ComK1 (Davids Biotechnologie) for 2h at 4°C with shaking and then incubated with 1:25000 goat anti-Rabbit IgG [HRP] (0.8 mg/mL; Davids Biotechnologie) for 2h at 4°C with shaking. After several washes, revelation was done using the Pierce^TM^ ECL Westen Blotting Substrate kit (ThermoFisher Scientific) and the membrane was scanned using a ChemiDoc^TM^ MP imaging system (BIO-RAD).

### RNA-sequencing. Sampling and isolation

Four cultures (biological replicates) of a wild type *S. aureus* strain were grown in CS2 medium at 37 °C and 180 rpm. Samples were taken at three different time points corresponding to different OD and O2 concentrations (Time1, OD= 0.08, O2= 12%; Time2, OD= 0.11, O2=0.05%; Time3, OD= 0.17, O2=0.05%; see arrows in **Fig. 5a**). It is important to note that the OD values shown here correspond to the height of 50 μL of culture in a 96-well plate rather than the typical 1 cm pathlength (for reference see (22)). To quench cellular metabolism / transcription and to stabilize RNAs, cells were harvested by centrifugation at 10,000g for 1 min at 4 °C and the pellets immediately frozen in liquid nitrogen before storage at −80 °C. For extraction of RNA, cells were lysed using Lysing Matrix B and a FastPrep instrument (both MP Biomedicals), and RNA were isolated using the RNeasy Mini Kit (Qiagen). RNAs were treated with TURBO DNase (Ambion), purified using the RNA Cleanup protocol from the RNeasy Mini Kit (Qiagen), and stored at −80 °C. The integrity of the RNA was finally analyzed using an Agilent Bioanalyzer (Agilent Technologies).

rRNA depletion, library construction and sequencing. Removal of 23 S, 16 S, and 5 S rRNA using the RiboZero rRNA Removal Kit (Epicenter) (two times), strand specific library construction yielding fragments of size range 100–500 bp, pooling of the 12 indexed libraries, sequencing in one flow-cell lane on a Illumina HiSeq2000 instrument with a 75 nt paired end protocol, and demultiplexing of the 12 samples of indexed reads was performed by the “Next Generation Sequencing (NGS) Core Facility” from the Institute for the Integrative Biology of the Cell (I2BC, Gif sur Yvette, France).

Read mapping and analysis of differential expression. A transcriptomic analysis was done using DicoExpress, a script-based tool allowing neutral comparisons studies (34). First, a filtering step was done to discard the low mean normalized counts using a CPM cut-off of 1. Data were normalized using the Trimmed Mean of M-values (TMM) method to homogenize the library size between samples (**Fig. Supp. Fig. 8a**). In the same way, the means of the gene expression for each sample were equalized (**Supp. Fig. 8b**). A Principal Component Analysis (PCA, Supp. Fig. 8c) and a Euclidean distance heatmap (**Supp. Fig. 8d**) of the dataset were also done on normalized counts to ensure the relevance of the dataset and to demonstrate the robustness of 613 the quality controls performed on the raw data. Finally, a differential analysis (between Time 1 (n = 4), Time 2 (n = 4) and Time 3 (n = 4)) was performed using the GLM_Contrasts function (GLM: generalized linear model) function from the edgeR package. To establish the list of the differentially expressed genes (DEGs), we kept the 808 genes with a p-value<0.01 and an absolute log2(FC)>1. From this list, 6 gene clusters were established using the coseq function, settled with recommended parameters. Each cluster is characterized by a specific expression pattern (increase or decrease between each time point).

## Data accessibility

The whole set of RNA-seq data is compiled and accessible under the GEO submission GSE281887.

## Reproducibility and statistical analysis

Expression profile. Every expression profile is represented as the mean of the transcription rate from 5 independent biological replicates (with at least 3 technical replicates each). Error bars are confidence intervals based on a Student Test (p-value<0.05; n=5).

Growth and growth rate curves. Growth and growth rate curves are shown as the mean of 15 replicates (5 independent biological replicates with at least 3 technical replicates each).

### Western blot

The western blot experiment shown in Fig. 3b has been repeated 3 times.

RNA-seq. The differential analysis (between Time 1 (n = 4), Time 2 (n = 4) and Time 3 (n = 4)) was performed using the GLM_Contrasts function (GLM: generalized linear model) function from the edgeR package. To establish the list of the differentially expressed genes (DEGs), we kept the 808 genes with a p-value<0.01 and an absolute log2(FC)>1. From this list, 6 gene clusters were established using the coseq function, settled with recommended parameters.

“n” represents the number of experiments considered to realize the statistical analysis.

## Acknowledgments

This work was supported by a “Young Researcher grant” from the French National Research Agency to Nicolas Mirouze (ANR-18-CE35-0004 GenTranSa) and a seed funding from the Interdisciplinary object MICROBES (“PROGRESS” project) from the Paris-Saclay University.

This work was supported by the “Fondation pour la Recherche Médicale” (FRM, grant number ECO202306017342), to Pierre Poirette.

This work was supported by a Chinese Scholarship Council (CSC) PhD grant awarded to Shi Yuan Feng. All RNA-sequencing experiments were performed in collaboration with the Next Generation Sequencing facility, I2BC in Gif-sur-Yvette – France.

Lastly, we wanted to thank Elodie Marchadier for her help with the analysis of the differentially expressed genes and gene set co-regulation study.

## Author contributions

Conceptualization, N.M.; Methodology, N.M., P.P, Y.H, S.Q.C. and S.M.; Investigation, P.P, S.Y.F, Y.H., Y.A., S.Q.C, S.M., M.R., M.D., R.M., T.H. and P.B.; Writing original draft, N.M., P.P.; Funding acquisition, N.M., P.P. and S.Y.F.; Resources, N.M.; Supervision, N.M., Y.H., S.Q.C. and S.M.

## References

1. Johnston C, Martin B, Fichant G, Polard P, Claverys JP. Bacterial transformation: distribution, shared mechanisms and divergent control. Nat Rev Microbiol. 2014;12(3):181–96. doi:10.1038/nrmicro3199 PubMed PMID: 24509783.

2. Griffith F. The Significance of Pneumococcal Types. J Hyg (Lond). 1928/01/01. 1928;27(2):113–59. PubMed PMID: 20474956.

3. Claverys JP, Martin B, Polard P. The genetic transformation machinery: composition, localization, and mechanism. FEMS Microbiol Rev. 2009/02/21. 2009;33(3):643–56. doi:10.1111/j.1574-6976.2009.00164.x PubMed PMID: 19228200.

4. Croucher NJ, Harris SR, Fraser C, Quail MA, Burton J, van der Linden M, et al. Rapid Pneumococcal Evolution in Response to Clinical Interventions. Science (1979). 2011 Jan 28;331(6016):430–4. doi:10.1126/science.1198545

5. Claverys JPP, Prudhomme M, Martin B. Induction of Competence Regulons as a General Response to Stress in Gram-Positive Bacteria. Annu Rev Microbiol. 2006/06/15. 2006 Oct 1;60(1):451–75. doi:10.1146/annurev.micro.60.080805.142139 PubMed PMID: 16771651.

6. Seitz P, Blokesch M. Cues and regulatory pathways involved in natural competence and transformation in pathogenic and environmental Gram-negative bacteria. FEMS Microbiology Reviews. 2013. p. 336–63. doi:10.1111/j.1574-6976.2012.00353.x PubMed PMID: 22928673.

7. Süel GM, Garcia-Ojalvo J, Liberman LM, Elowitz MB. An excitable gene regulatory circuit induces transient cellular differentiation. Nature. 2006 Mar;440(7083):545–50. doi:10.1038/nature04588

8. Ogura M, Yamaguchi H, Kobayashi K, Ogasawara N, Fujita Y, Tanaka T. Whole- genome analysis of genes regulated by the Bacillus subtilis competence transcription factor ComK. J Bacteriol. 2002/04/12. 2002;184(9):2344–51. PubMed PMID: 11948146.

9. Berka RM, Hahn J, Albano M, Draskovic I, Persuh M, Cui X, et al. Microarray analysis of the Bacillus subtilis K-state: genome-wide expression changes dependent on ComK. Mol Microbiol. 2002/03/29. 2002;43(5):1331–45. PubMed PMID: 11918817.

10. Dagkessamanskaia A, Moscoso M, Henard V, Guiral S, Overweg K, Reuter M, et al. Interconnection of competence, stress and CiaR regulons in Streptococcus pneumoniae: competence triggers stationary phase autolysis of ciaR mutant cells. Mol Microbiol. 2004/02/07. 2004;51(4):1071–86. PubMed PMID: 14763981.

11. Peterson S, Cline RT, Tettelin H, Sharov V, Morrison DA. Gene expression analysis of the Streptococcus pneumoniae competence regulons by use of DNA microarrays. J Bacteriol. 2000;182(21). doi:10.1128/JB.182.21.6192-6202.2000

12. Peterson SN, Sung CK, Cline R, Desai B V., Snesrud EC, Luo P, et al. Identification of competence pheromone responsive genes in Streptococcus pneumoniae by use of DNA microarrays. Mol Microbiol. 2004 Feb;51(4):1051–70. doi:10.1046/j.1365-2958.2003.03907.x

13. Chen JD, Morrison DA. Modulation of competence for genetic transformation in Streptococcus pneumoniae. J Gen Microbiol. 1987;133(7). doi:10.1099/00221287-133-7-1959

14. Hamoen LW. Controlling competence in Bacillus subtilis: shared use of regulators. Microbiology (N Y). 2003 Jan 1;149(1):9–17. doi:10.1099/mic.0.26003-0

15. Prudhomme M, Attaiech L, Sanchez G, Martin B, Claverys JP. Antibiotic stress induces genetic transformability in the human pathogen Streptococcus pneumoniae. Science (1979). 2006/07/11. 2006;313(5783):89–92. doi:10.1126/science.1127912 PubMed PMID: 16825569.

16. Maziero M, Lane D, Polard P, Bergé M. Fever-like temperature bursts promote competence development via an HtrA-dependent pathway in Streptococcus pneumoniae. PLoS Genet. 2023 Sep 12;19(9):e1010946. doi:10.1371/journal.pgen.1010946

17. Lee MS, Morrison DA. Identification of a new regulator in Streptococcus pneumoniae linking quorum sensing to competence for genetic transformation. J Bacteriol. 1999;181(16):5004–16. PubMed PMID: 10438773.

18. Morikawa K, Takemura AJ, Inose Y, Tsai M, Nguyen Thi le T, Ohta T, et al. Expression of a cryptic secondary sigma factor gene unveils natural competence for DNA transformation in Staphylococcus aureus. PLoS Pathog. 2012/11/08. 2012;8(11):e1003003. doi:10.1371/journal.ppat.1003003 PubMed PMID: 23133387.

19. Morikawa K, Inose Y, Okamura H, Maruyama A, Hayashi H, Takeyasu K, et al. A new staphylococcal sigma factor in the conserved gene cassette: functional significance and implication for the evolutionary processes. Genes to Cells. 2003 Aug;8(8):699–712. doi:10.1046/j.1365-2443.2003.00668.x

20. Fagerlund A, Granum PE, Havarstein LS. Staphylococcus aureus competence genes: mapping of the SigH, ComK1 and ComK2 regulons by transcriptome sequencing. Mol Microbiol. 2014;94(3):557–79. doi:10.1111/mmi.12767 PubMed PMID: 25155269.

21. van Sinderen D, Luttinger A, Kong L, Dubnau D, Venema G, Hamoen L. comK encodes the competence transcription factor, the key regulatory protein for competence development in Bacillus subtilis. Mol Microbiol. 1995;15(3):455–62. PubMed PMID: 7783616.

22. Feng SY, Hauck Y, Morgene F, Mohammedi R, Mirouze N. The complex regulation of competence in Staphylococcus aureus under microaerobic conditions. Commun Biol. 2023 May 12;6(1):512. doi:10.1038/s42003-023-04892-1

23. Martin B, Garcia P, Castanie MP, Claverys JP. The recA gene of Streptococcus pneumoniae is part of a competence-induced operon and controls lysogenic induction. Mol Microbiol. 1995/01/01. 1995;15(2):367–79. PubMed PMID: 7538190.

24. Moriyama EH, Niedre MJ, Jarvi MT, Mocanu JD, Moriyama Y, Subarsky P, et al. The influence of hypoxia on bioluminescence in luciferase-transfected gliosarcoma tumor cells in vitro. Photochemical and Photobiological Sciences. 2008;7(6). doi:10.1039/b719231b

25. Feng SY, Arab Y, Hauck Y, Poirette P, Noiray M, Quevillon-Cheruel S, et al. ComK2 represses competence development for natural transformation in Staphylococcus aureus grown under strong oxygen limitation. Commun Biol. 2025 Oct 2;8(1):1416. doi:10.1038/s42003-025-08816-z

26. Ibarra JA, Pérez-Rueda E, Carroll RK, Shaw LN. Global analysis of transcriptional regulators in Staphylococcus aureus. BMC Genomics. 2013;14(1):126. doi:10.1186/1471-2164-14-126

27. Sonenshein AL. CodY, a global regulator of stationary phase and virulence in Gram-positive bacteria. Curr Opin Microbiol. 2005 Apr;8(2):203–7. doi:10.1016/j.mib.2005.01.001

28. Serror P, Sonenshein AL. CodY is required for nutritional repression of Bacillus subtilis genetic competence. J Bacteriol. 1996 Oct;178(20):5910–5. doi:10.1128/jb.178.20.5910-5915.1996

29. Mirouze NN; FC; CLR, Ferret C, Cornilleau C, Carballido-López R. Antibiotic sensitivity reveals that wall teichoic acids mediate DNA binding during competence in Bacillus subtilis. Nat Commun. 2018 Dec 29;9(1):5072. doi:10.1038/s41467-018-07553-8

30. Lo Scrudato M, Blokesch M. A transcriptional regulator linking quorum sensing and chitin induction to render Vibrio cholerae naturally transformable. Nucleic Acids Res. 2013 Apr;41(6):3644–58. doi:10.1093/nar/gkt041

31. Toussaint F, Henry de Frahan M, Poncelet F, Ladrière JM, Horvath P, Fremaux C, et al. Unveiling the regulatory network controlling natural transformation in lactococci. PLoS Genet. 2024 Jul 1;20(7):e1011340. doi:10.1371/journal.pgen.1011340

32. Mirouze N, Prepiak P, Dubnau D. Fluctuations in spo0A transcription control rare developmental transitions in bacillus subtilis. PLoS Genet. 2011;7(4). doi:10.1371/journal.pgen.1002048

33. Monk IR, Shah IM, Xu M, Tan MW, Foster TJ. Transforming the Untransformable: Application of Direct Transformation To Manipulate Genetically Staphylococcus aureus and Staphylococcus epidermidis. Novick RP, editor. mBio. 2012 May 2;3(2). doi:10.1128/mBio.00277-11

34. Lambert I, Paysant-Le Roux C, Colella S, Martin-Magniette ML. DiCoExpress: a tool to process multifactorial RNAseq experiments from quality controls to co-expression analysis through differential analysis based on contrasts inside GLM models. Plant Methods. 2020 Dec 12;16(1):68. doi:10.1186/s13007-020-00611-7

